# Genomic and functional characterization of *Bacillus* sp. strain Bva_UNVM-123 reveals resistance and environmental adaptation traits

**DOI:** 10.64898/2026.03.31.715617

**Authors:** Cecilia Peralta, Diego Herman Sauka, Verónica Felipe, Sergio Redondo-Moreno, Eleodoro E. Del Valle, Leopoldo Palma

## Abstract

The *Bacillus* genus comprises physiologically versatile, endospore-forming bacteria widely distributed in natural environments. In this study, we report the isolation and genomic characterization of strain Bva_UNVM-123, recovered from agricultural soil in Pergamino, Argentina. Whole-genome sequencing using Illumina technology yielded a 5.1 Mbp draft genome assembled in 67 contigs with a GC content of 36%. Comparative genomic analyses using the TYGS server and digital DNA–DNA hybridization (dDDH) values supported its classification as a potentially novel species within the *Bacillus* sensu lato (s.l.) group. Genome annotation revealed 4,866 protein-coding genes, including multiple determinants conferring resistance to antibiotics (e.g., fosfomycin, tetracycline, beta-lactams) and toxic heavy metals (e.g., arsenic, cadmium, mercury), supporting its potential application in bioremediation. Additionally, PathogenFinder predicted a low probability of human pathogenicity (0.207), reinforcing its safety for environmental use. Functional classification based on Swiss-Prot further supported a metabolically versatile profile and revealed the presence of resistance-related categories associated with environmental adaptation. This study adds to the growing knowledge of environmental *Bacillus* species and their biotechnological potential.

## Introduction

Species belonging to the *Bacillus* genus are known for their remarkable natural properties, which range from the synthesis of various toxins (e.g., cereulide) to the production of proteins and enzymes with numerous biotechnological applications, including fermentation, biodegradation, and the production of secondary metabolites. Many of these organisms can synthesize a wide range of bioactive compounds, including antibiotics (e.g., bacitracina), enzymes (amylases, proteases, lipases, etc.), and biopolymers, which can be used in industrial applications (Herrmann et al., 2024).

The genes responsible for these biosynthetic pathways are often located on extrachromosomal DNA elements, such as plasmids or bacteriophages, which can harbor additional virulence factors including toxins and antibiotic resistance genes (Bennett, 2008).

These plasmids are typically transferable, and through horizontal gene transfer (HGT), they can be transmitted to related bacterial species (Sauka et al., 2024). Horizontal transfer of genetic material between bacteria plays a crucial role in the spread of antibiotic resistance, as it allows for rapid dissemination of resistance traits across different bacterial populations (Kaplan, 2014; Partridge et al., 2018). This transfer can occur between pathogenic and non-pathogenic bacteria, and in the case of non-pathogenic species, it can result in the presence of antibiotic resistance genes even in bacteria that do not cause disease (Martínez, 2008). Notably, recent studies emphasize the role of non-pathogenic soil bacteria in harboring mobile resistance elements that may be horizontally transferred to clinical pathogens under environmental stress conditions. Such non-pathogenic bacteria, isolated from various ecosystems like soil, water, or animal manure, are of great interest because they can serve as reservoirs for resistance genes that may eventually be transferred to pathogenic organisms under selective pressure (Larsson and Flach, 2022).

Antibiotic resistance genes are essential mechanisms used by pathogenic bacteria to evade the effects of antibiotics, a challenge that has become a major global health concern. However, they are not only found in pathogenic bacteria but are also commonly detected in non-pathogenic environmental microorganisms, including species used in biotechnology and environmental applications (Amos, 2013). This characteristic is particularly interesting from an environmental management perspective because non-pathogenic, antibiotic-resistant bacteria could potentially be utilized in bioremediation strategies. For example, certain *Bacillus* strains with intrinsic resistance to antibiotics can be employed in bioremediation efforts to clean up environments contaminated by antibiotics, particularly those originating from hospitals, dairy farms, and livestock operations, where residues of antibiotics often persist in soil and water (Gul et al., 2022). This provides a valuable opportunity for using these bacteria in the development of new biotechnological tools aimed at mitigating the environmental impact of antibiotic contamination, which is a growing concern in public health and environmental science (Apreja et al., 2022).

In addition to antibiotic resistance, several *Bacillus* species are known to tolerate or resist toxic heavy metals such as arsenic, cadmium, lead, copper, and mercury. This resistance is often mediated by efflux systems, metal-binding proteins, and enzymatic detoxification pathways encoded in their genomes (Margaryan et al., 2021; Prabhakaran et al., 2016). These properties make them promising candidates for bioremediation strategies aimed at decontaminating polluted environments. The capacity to thrive in metal-contaminated soils, together with their ability to form resilient endospores, enhances their potential in long-term or large-scale remediation projects (Singh, 2014).

The capacity of *Bacillus* species to adapt to diverse environments, combined with their resistance to antibiotics and heavy metals, makes them ideal candidates for further research in the field of bioremediation. By leveraging their natural resistance mechanisms, it may be possible to engineer more efficient systems for cleaning up contaminated environments, particularly those impacted by human activity.

In this work, we aimed to sequence and analyze the genome of strain Bva_UNVM-123, a newly isolated *Bacillus* strain from soil, to explore its taxonomic identity and biotechnological potential. The assembled draft genome sequence of Bva_UNVM-123 provides insights into its genomic makeup, revealing potential resistance genes against antibiotics and heavy metals, which may contribute to its potential as a bioremediation agent. This study contributes to a deeper understanding of the genetic factors underlying resistance in *Bacillus* species, further supporting their application in environmental remediation. Additionally, genome-based analyses suggest that strain Bva_UNVM-123 may represent a novel species within the *Bacillus* sensu lato (s.l.) group, for which the name *Bacillus pergaminensis* has been proposed.

## Materials and Methods

### Isolation and Morphological Characterization

The strain Bva_UNVM-123 was isolated from a soil sample collected from soybean field in Pergamino, Buenos Aires Province (Argentina), as part of a screening program aimed at identifying entomopathogenic bacteria (e.g., *Bacillus thuringiensis*) as previously described (Iriarte et al., 1998).

Those strains showing a *B. thuringiensis*-like (matte white and with uneven borders) phenotype were reisolated in axenic cultures and observed by phase contrast microscopy (Nikon Eclipse Ti, Japan). Gram staining was performed using a commercial kit (Biopack, Buenos Aires, Argentina), following the manufacturer’s instructions. In addition, the strain was also examined using Scanning Electron Microscopy (SEM) at the Comprehensive Center for Electron Microscopy (CONICET-Universidad Nacional de Tucumán, Argentina). The strain is maintained as glycerol suspension (50% v/v) at −80 °C.

The type strain Bva_UNVM-123^t^ was deposited in the International Culture Collection of the Institute of Microbiology and Agricultural Zoology at INTA Castelar (IMYZA-INTA) (access number pending).

### Biochemical Characterization and Antibiotic Resistance Analyses

NaCl tolerance was evaluated by culturing the strain on TSA medium supplemented with NaCl concentrations ranging from 2.5% to 7.5% (w/v) and incubating for 72 hours. Anaerobic growth was assessed using the candle jar technique. Motility was tested on semisolid agar medium (0.75% w/v). Hemolytic activity was tested on blood agar plates supplemented with 5% sheep blood (Britania Laboratories, Argentina) incubated at 37 °C for 48 hours.

The sensitivity of strain Bva_UNVM-123 to various antibiotics, chloramphenicol (30 µg), novobiocin (5 µg), ampicillin (10 µg), tetracycline (30 µg), and penicillin (10 U) (Britania Laboratories, Argentina), was assessed on Mueller–Hinton agar using the disc diffusion method. Oxidase and catalase activities were determined using standard microbiological procedures. Additional physiological and biochemical characterizations were conducted using the API 50CHB, API ZYM, and API 20A systems (bioMérieux), following the protocols provided by the manufacturer.

Fatty acid profiling of the strain was performed using the MIDI Sherlock Microbial Identification System (version 6.2B) following the methodology described by Ferrari et al., (2018). The sample was analyzed using the RTSBA6 method, and fatty acids were identified based on their equivalent chain length (ECL) and retention time (Ferrari et al., 2018).

### DNA isolation, Genome assembly and Annotation and Species Delimitation

For DNA extraction, strain Bva_UNVM-123 was grown in LB broth for 16 hours. Genomic DNA, including chromosomal and plasmid DNA, was extracted using the Wizard Genomic DNA Purification Kit (Promega, Madison, WI, USA), following the manufacturer’s protocol for gram-positive bacteria. The extracted DNA was analyzed by electrophoresis in 1% agarose gels, stained with SYBR Safe (ThermoFisher Scientific, Waltham, MA, USA), and quantified using a PICODROP PICO 100 μL spectrophotometer.

The extracted DNA was subjected to PCR to confirm its belonging to the *Bacillus* genus (Wu et al., 2006). The PCR assay was performed using the methodology described by Wu et al. (2006), which involves a group-specific primer set designed for *Bacillus* species identification.

Then, the purified DNA was used to construct a pooled Illumina sequencing library, which was sequenced at the Genomics Unit of the National Institute of Agricultural Technology (INTA, Argentina) using Illumina high-throughput sequencing technology. Raw sequencing reads were processed to remove low-quality regions and assembled into contigs using Velvet within Geneious version R11 software (www.geneious.com) with default parameters for *de novo* assembly.

Assembly statistics were evaluated using QUAST v4.5 (Gurevich et al., 2013), and genome completeness and contamination were assessed using CheckM (Parks et al., 2015). Gene predictions and functional annotations were done via PGAP [16] and validated through the RAST server (Aziz et al., 2008).

Genome annotation was performed using the NCBI Prokaryotic Genome Annotation Pipeline (2023 release) and the RAST server (Aziz et al., 2008).

Species delimitation was carried out using the Type (Strain) Genome Server (Meier-Kolthoff and Goker, 2019) and PubMLST (Jolley et al., 2018). Secondary metabolites searches were performed using antiSmash server (Blin et al., 2019). Average Nucleotide Identity (ANI) values were also calculated using FastANI (Jain et al., 2018) by comparing the genome of strain Bva_UNVM-123 against closely related taxa identified through TYGS analysis, including representatives of the genera *Lysinibacillus, Cytobacillus, Neobacillus*, and *Ferdinandcohnia*.

### Functional classification based on Swiss-Prot homology

RAST server predicted proteins were additionally queried against a local UniProtKB/Swiss-Prot database (Boutet et al., 2007) using BLASTp (Altschul et al., 1990) with an e-value cutoff of 1e-10, retaining only the top-scoring hit for each query. Significant hits were classified into broad functional groups according to keyword-based annotation mining, and category frequencies were plotted using the ggplot2 package in R (Wickham, 2016).

## Results

### Morphological Characterization

The strain formed opaque colonies with a matte texture and creamy white color after overnight growth on TSA medium at 37 °C during 72 h. Microscopic examination revealed rod-shaped vegetative cells with terminal deforming spores (Figure 1A and B). The terminal positioning of the endospore is consistent with known morphological traits in environmental *Bacillus* species (Vos et al., 2011).

**Figure 1.**
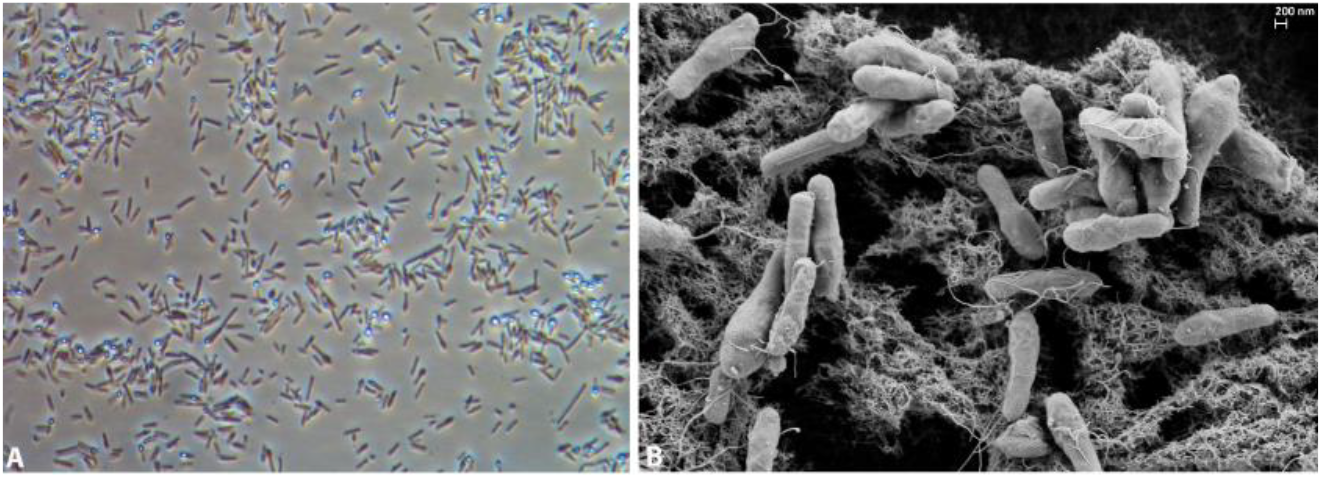
Microscopic analysis of strain Bva_UNVM-123 grown on TSA medium (72 h/37 °C). A) Morphological characterization using phase-contrast microscopy and B) Scanning electron microscopy. A scale bar (200 nm) was included in SEM images to reflect structural dimensions.

### Biochemical Characterization

Cells are Gram-stain-positive, facultatively anaerobic, non-motile, and form terminal endospores. The strain was able to grow on nutrient agar (NA), casein–collagen–yeast extract agar (CCY), tryptic soy agar (TSA), and brain heart infusion agar. Optimal growth was observed on TSA at 37 °C and the strain was non-hemolytic. The strain tested positive for both catalase and oxidase activities and when cultured in the presence of chloramphenicol, did not exhibit measurable inhibition zones.

Additional physiological and enzymatic tests were carried out using the API 50CHB, API ZYM, and API 20A systems. However, the results obtained were inconclusive due to limited growth under assay conditions, including incubation at 30 °C and 37 °C.

The fatty acid profile of strain Bva_UNVM-123 consisted primarily of branched-chain saturated fatty acids, a chemotaxonomic trait typical of gram-positive bacteria, particularly members of the genus *Bacillus* or related taxa. The major components included iso- and anteiso-fatty acids such as iso-C15:0, anteiso-C15:0, iso-C16:0, and iso-C17:0. A total of 61.21% of the detected fatty acids were identified, with iso-C15:0 representing the most abundant component (32.46%). It is worth noting that obtaining the fatty acid profile was technically challenging due to the limited growth of the strain under standard laboratory conditions (e.g., LB medium), which may have affected biomass yield and detection sensitivity. The complete fatty acid profile is presented in Table 1.

**Table 1.** Key Major fatty acid components of strain Bva_UNVM-123. Only those representing ≥3% of the total identified fatty acids are shown. Summed Feature 4 includes fatty acids that could not be resolved individually by the system.

| Fatty Acid | Composition (%) | Other Characteristics |
| --- | --- | --- |
| iso-C15:0 | 32.46 | Most abundant component |
| anteiso-C15:0 | 7.28 | - |
| iso-C16:0 | 5.21 | - |
| C16:0 | 4.39 | Also known as palmitic acid |
| iso-C17:0 | 6.16 | - |
| anteiso-C17:0 | 3.69 | - |
| Summed Feature 4 | 5.20 | Includes C17:1 iso I / anteiso B isomers |

### Genome Characterization and Species Identification

The Illumina sequencing produced 14,532,725 high-quality paired end reads, resulting in 67 assembled contigs with a total genome size of 5,119,019 bp in size and 36% G+C content (Table 2).

**Table 2.** Key features statistics and quality analysis of the draft genome assembly of Bva_UNVM-123 strain obtained with QUAST, CheckM and the NCBI Prokaryotic Genome Annotation Pipeline (PGAP).

| Feature statistics | Value |
| --- | --- |
| Assembled Genome Size (bp) | 5,119,099 |
| Contig Number | 67 |
| Largest contig (bp) | 583,230 |
| N50 (bp) | 341,242 |
| N's | 5074 |
| GC content (%) | 36 |
| Suspected Contamination (%)* | 3.85 |
| Estimated Completeness (%)* | 95.85 |
| Estimated Coverage (×) | 400 |
| PGAP Predicted Genes | 5139 |
| PGAP Predicted Protein Coding Genes | 4866 |

\*CheckM determinations.

Table 3 highlights multiple genes potentially related to antibiotic resistance, identified using the RAST annotation server. The presence of genes conferring resistance to fosfomycin (fosB), tetracyclines, chloramphenicol, fosmidomycin, kanamycin, oxetanocin A, and various beta-lactams (e.g., beta-lactamase class A-like and penicillin-binding proteins) suggests that strain Bva_UNVM-123 possesses a broad resistance profile.

**Table 3.**
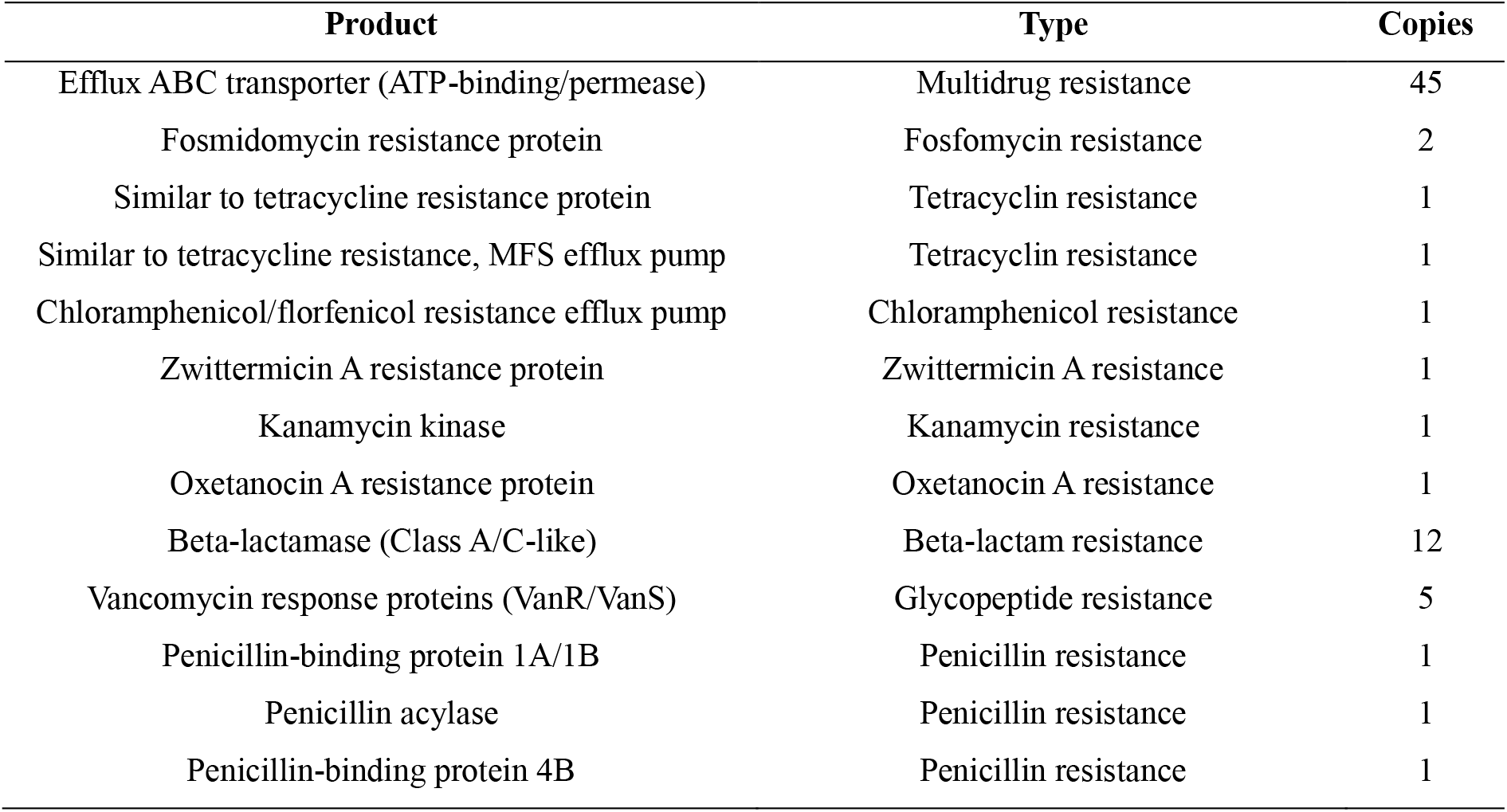
Putative annotated genes related to antibiotic resistance found at the genome sequence of Bva_UNVM-123 strain using RAST server.

| Product | Type | Copies |
| --- | --- | --- |
| Efflux ABC transporter (ATP-binding/permease) | Multidrug resistance | 45 |
| Fosmidomycin resistance protein | Fosfomycin resistance | 2 |
| Similar to tetracycline resistance protein | Tetracyclin resistance | 1 |
| Similar to tetracycline resistance, MFS efflux pump | Tetracyclin resistance | 1 |
| Chloramphenicol/florfenicol resistance efflux pump | Chloramphenicol resistance | 1 |
| Zwittermicin A resistance protein | Zwittermicin A resistance | 1 |
| Kanamycin kinase | Kanamycin resistance | 1 |
| Oxetanocin A resistance protein | Oxetanocin A resistance | 1 |
| Beta-lactamase (Class A/C-like) | Beta-lactam resistance | 12 |
| Vancomycin response proteins (VanR/VanS) | Glycopeptide resistance | 5 |
| Penicillin-binding protein 1A/1B | Penicillin resistance | 1 |
| Penicillin acylase | Penicillin resistance | 1 |
| Penicillin-binding protein 4B | Penicillin resistance | 1 |

Table 4 lists genes potentially associated with resistance to toxic heavy metals, including arsenic, cadmium, zinc, cobalt, lead, mercury, and tellurite. These genes encode various efflux pumps, ATPases, and regulatory proteins, such as TehB, ArsD, ACR3, and CzcD, which are known to facilitate resistance and detoxification (Bruins et al., 2000; Nies, 2003).

**Table 4.** Putative heavy-metal resistance genes found at the genome sequence of Bva_UNVM-123 strain using RAST server.

| Product | Metal | Copies |
| --- | --- | --- |
| Tellurium resistance proteins (TerD, TerB) | Tellurium | 5 |
| Arsenical resistance proteins (Acr3, ArsR, ATPase) | Arsenic | 9 |
| Cadmium/Zinc/Cobalt transporters (P-type ATPase, CzcD) | Cd/Zn/Co | 7 |
| Copper chaperones and transporters (CopZ, CopA) | Copper | 4 |
| Zinc ABC transporter systems (ZnuA/B/C, Zur) | Zinc | 5 |
| Manganese efflux pump (MntP) | Manganese | 1 |
| Lead, Mercury, and other heavy-metal transporters | Pb/Hg/Cd/Zn | 6 |
| Multicopper oxidases and polyphenol oxidases | Copper | 1 |
| Aluminum resistance protein | Aluminum | 1 |

The Swiss-Prot-based functional classification revealed a proteome dominated by housekeeping and metabolic functions, together with a relevant proportion of transport-associated proteins and a smaller but clearly detectable set of resistance-related proteins. In particular, the identification of both antibiotic- and heavy metal-associated categories agrees with the resistance determinants detected by RAST and reinforces the proposed ecological adaptation and bioremediation potential of strain Bva_UNVM-123.

Metabolism represented the largest functional category (30.4%), followed by other/poorly resolved functions (30.3%), unknown proteins (12.2%), information processing (10.5%), and transport-related proteins (8.6%). Resistance-related categories, including heavy metal resistance (1.2%) and antibiotic resistance (0.4%), were also detected (Figure 2).

**Figure 2.**
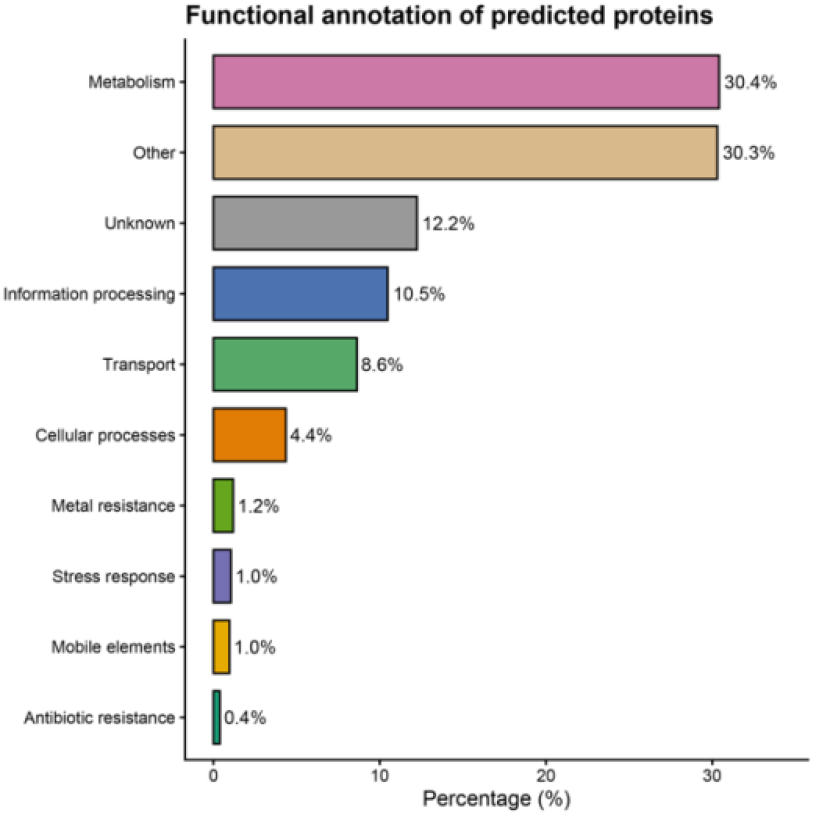
Functional classification of predicted proteins of strain Bva_UNVM-123 based on BLASTp searches against the Swiss-Prot database. Proteins were grouped into broad functional categories based on annotation keywords. Percentages are calculated relative to the total number of proteins with significant Swiss-Prot matches.

It should be noted that Swiss-Prot represents a curated but limited database, and therefore the proportion of annotated proteins reflects high-confidence functional assignments rather than the total functional potential of the genome.

Concerning species identification, preliminary PCR analysis showed that isolate 123 belongs to the genus *Bacillus* (Figure 3).

**Figure 3.**
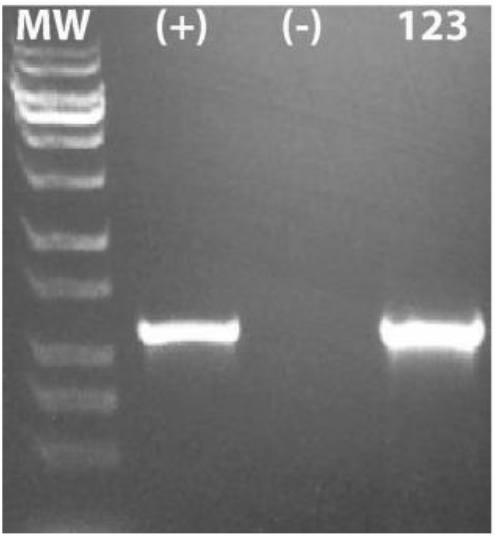
PCR analysis of strain Bva_UNVM-123 for the Bacillus genus showing a positive PCR amplicon (positive control: *B. thuringiensis* HD1, 123: Bva_UNVM-123 strain).

The genome-based phylogenetic tree revealed that strain Bva_UNVM-123 clusters within a lineage comprising members of the *Bacillus* sensu lato (s.l.) group, showing closest affiliation to *Lysinibacillus* and related genera (Figure 4). This placement is consistent with the complex and recently revised taxonomy of the *Bacillus* group and supports its classification as a distinct genomic lineage.

**Figure 4.**
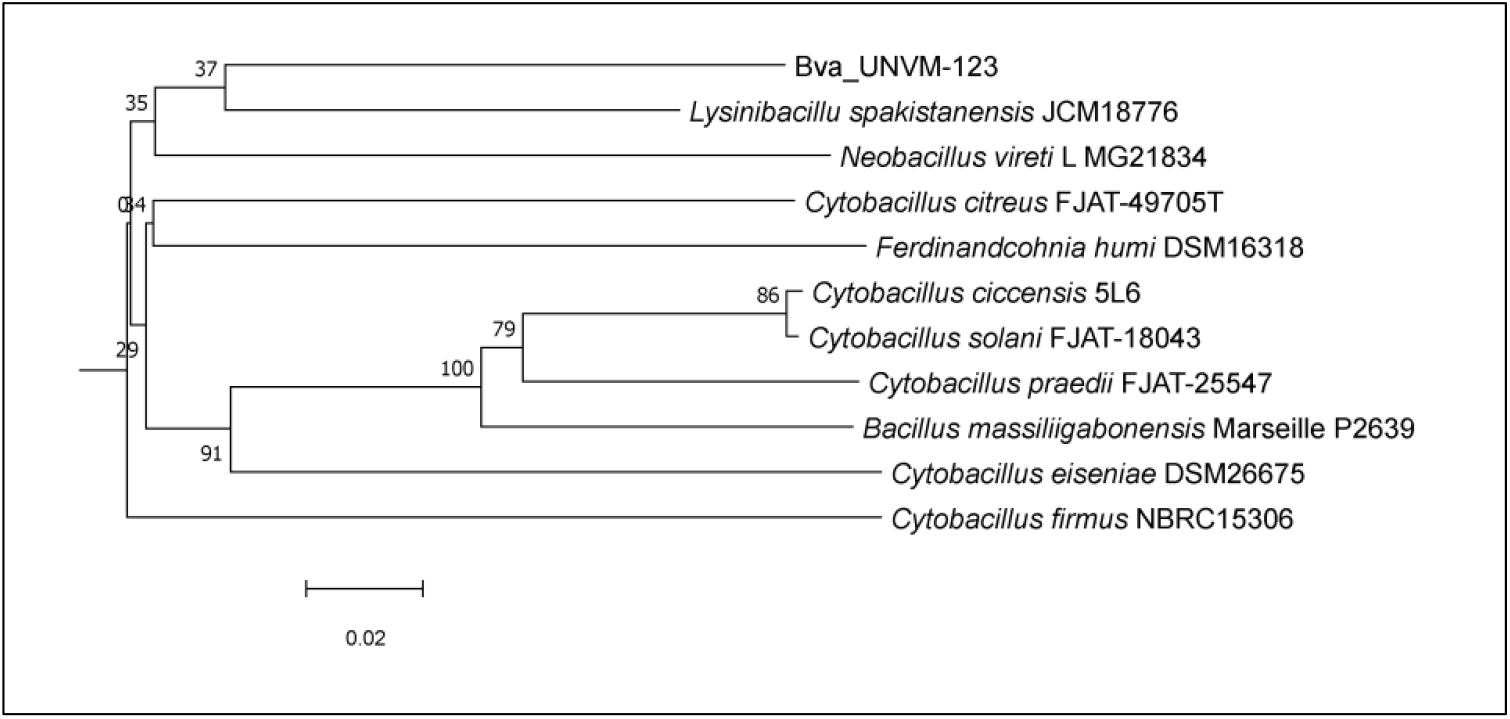
Genome-based phylogenetic tree inferred using the GBDP method via the TYGS server, showing the position of strain Bva_UNVM-123 among closely related taxa within the *Bacillus* sensu lato (s.l.) group. The strain clusters in proximity to members of the genera *Lysinibacillus, Cytobacillus*, and *Neobacillus*. Branch lengths are scaled in terms of GBDP distance, and bootstrap values (>60%) based on 100 pseudoreplicates are indicated at the nodes.

Moreover, average Nucleotide Identity (ANI) analysis revealed low similarity between strain Bva_UNVM-123 and the closest related genomes identified by TYGS, with all values well below the accepted species threshold of 95% (Table 5), confirming its distinct genomic status.

**Table 5.** Average Nucleotide Identity (ANI) values between strain Bva_UNVM-123 and closely related genomes identified by TYGS. ANI values correspond to the mean of reciprocal (bidirectional) comparisons calculated using FastANI.

| Reference genome | ANI (%) |
| --- | --- |
| *Cytobacillus solani* FJAT-18043 | 79.05 |
| *Cytobacillus solani* 5L6 | 78.87 |
| *Cytobacillus praedii* FJAT-25547 | 78.57 |
| *Cytobacillus massiliigabonensis* Marseille P2639 | 78.32 |
| *Cytobacillus citreus* FJAT-49705 | 78.18 |
| *Cytobacillus eiseniae* DSM26675 | 78.14 |
| *Neobacillus vireti* LMG21834 | 77.47 |
| *Cytobacillus firmus* NBRC15306 | 77.47 |
| *Lysinibacillus pakistanensis* JCM18776 | 77.42 |
| *Ferdinandcohnia humi* DSM16318 | 77.06 |

In silico sequence typing using the PubMLST database identified alleles for multiple loci (n = 53); however, the allelic profile did not match any previously defined sequence type, suggesting that strain Bva_UNVM-123 represents a novel sequence type. This result is consistent with the phylogenomic analyses and further supports the distinct genomic position of the strain.

Based on PathogenFinder 1.1, the strain was predicted as non-pathogenic, with a human pathogenicity probability of 0.207, indicating its safety for environmental applications (Cosentino et al., 2013).

## Discussion

After comprehensive genomic and phenotypic analyses, including phylogenomic reconstruction and digital DNA–DNA hybridization (dDDH) values below the 70% species threshold, strain Bva_UNVM-123 likely represents a novel species within the *Bacillus* sensu lato (s.l.) group. These findings suggest that it constitutes a distinct genomic lineage within this taxonomic framework, for which the name *Bacillus pergaminensis* has been proposed. This interpretation is further supported by ANI values below 80% against all closely related genomes tested, indicating a high level of genomic divergence from currently described taxa within the *Bacillus* sensu lato (s.l.) group. The low ANI values observed in this study may also indicate a deeper level of taxonomic divergence; however, additional analyses would be required to clarify its precise taxonomic placement.

The genome of strain Bva_UNVM-123 reveals multiple adaptive traits that support its role in bioremediation. In line with this, preliminary phenotypic assays showed that strain Bva_UNVM-123 exhibited no measurable inhibition zones against chloramphenicol and novobiocin, suggesting intrinsic tolerance that may correspond to the resistance genes found in the genome. Nonetheless, further MIC assays are necessary to validate this resistance. The relatively high proportion of uncharacterized proteins highlights the presence of potentially novel functions, which may represent unexplored metabolic or adaptive capabilities.

In addition, the presence of antibiotic resistance genes such as fosB, tetA, bla-like beta-lactamases, and multidrug efflux systems suggests long-term exposure to antibiotic residues in soil environments. This aligns with findings that environmental *Bacillus* species often act as reservoirs for antibiotic resistance genes due to selective pressure in natural and agricultural soils (Larsson and Flach, 2022; Martínez, 2008). This resistance could be determinant and particularly relevant for bioremediation applications, where bacteria must survive in antibiotic-contaminated environments. These results are consistent with the resistance determinants identified through RAST annotation (Tables 3 and 4), further supporting the functional potential of the strain for survival in contaminated environments.

The use of API-based systems for metabolic profiling did not yield conclusive results. This outcome is frequently observed with environmental isolates, which often exhibit poor growth or atypical metabolic patterns when tested in commercial platforms originally designed for clinical or fast-growing laboratory strains. These limitations highlight the need to optimize test conditions or employ alternative phenotypic strategies when characterizing environmentally adapted bacteria. For instance, even in industrial contexts such as dairy processing environments, phenotypic microbiological techniques have shown limited reliability for environmental strains like *Pseudomonas* spp. and *Bacillus* spp., often resulting in inconclusive species-level identifications (Wiedmann et al., 2000). Consequently, in this study, species identification relied primarily on genome-based approaches, which provided higher taxonomic resolution and reliability.

Additionally, the fatty acid profile of strain Bva_UNVM-123 did not match any entry in the RTSBA6 reference library, which is consistent with the genomic evidence suggesting novelty. However, the total response signal obtained in both replicates was below the recommended threshold for reliable identification. This limitation was most likely due to the poor growth of the strain under the conditions used for biomass collection, rather than to methodological issues. Further optimization of culture conditions is therefore needed to obtain sufficient biomass for accurate chemotaxonomic profiling.

Moreover, the strain harbors a diverse array of heavy metal resistance genes, including *arsD, acr3, czcD*, and *tehB*, which confer tolerance to arsenic, cadmium, mercury, and tellurite. These genes are commonly found in metal-contaminated environments and are considered key biomarkers for bacterial adaptation and survival in such niches (Bruins et al., 2000; Margaryan et al., 2021; Nies, 2003). The abundance and diversity of these genes suggest that strain Bva_UNVM-123 is well adapted to metal-contaminated environments. This further supports its potential use in bioremediation, particularly in soils impacted by industrial or agricultural pollutants. The presence of both antibiotic and heavy metal resistance-related functional categories, even at relatively low proportions, is particularly relevant, as these traits are often co-selected in environmental bacteria exposed to anthropogenic pressures. This supports the hypothesis that strain Bva_UNVM-123 may be adapted to environments impacted by agricultural and chemical inputs.

The non-pathogenic nature of the strain, as predicted by PathogenFinder, along with the absence of virulence determinants and hemolytic activity, strongly suggests its suitability for environmental application. This safety profile is crucial for any candidate microorganism intended for bioremediation purposes, as environmental deployment requires strict biosafety assessments (Cosentino et al., 2013).

Importantly, the genome of the strain Bva_UNVM-123 also provides a valuable resource for future biotechnological exploration. Its genetic repertoire could be exploited not only for bioremediation of antibiotic- and metal-polluted sites but also for the potential production of industrial enzymes or bioactive secondary metabolites, as often observed in *Bacillus* spp. (Blin et al., 2019; Herrmann et al., 2024).

## Conclusion

The genomic and functional characterization of strain Bva_UNVM-123 reveals a bacterium with a complex adaptive profile shaped by environmental pressures. Genome-based analyses, including TYGS phylogenomics and ANI, consistently indicate that this strain represents a highly divergent lineage within the *Bacillu*s sensu lato (s.l.) group, with ANI values below 80% against all closely related taxa. These results strongly support its classification as a previously uncharacterized species-level lineage.

The presence of a diverse repertoire of genes associated with resistance to antibiotics and heavy metals, together with transport systems and stress-response mechanisms, highlights the ecological versatility of this strain and its potential adaptation to contaminated environments. Functional classification further supports a metabolically active organism capable of coping with multiple environmental stressors.

Importantly, the predicted non-pathogenic profile and the absence of virulence-associated traits reinforce its suitability for environmental applications. Altogether, these findings position strain Bva_UNVM-123 as a promising candidate for future biotechnological exploitation, particularly in the context of bioremediation of environments impacted by anthropogenic pollutants.

This study contributes to expanding the current understanding of genomic diversity within the *Bacillus* sensu lato group and underscores the value of integrating genomic and functional approaches to uncover novel environmentally relevant bacterial lineages.

## Author contributions

Conceptualization, L.P., methodology, D.H.S, C.P and L.P.; formal analysis, D.H.S, C.P, V.F. and L.P.; investigation, D.H.S; C.P, V.F., E.E.D.V and L.P.; resources, L.P.; data curation, L.P.; writing—original draft preparation, C.P., L.P.; writing, review and editing, L.P. and D.H.S; visualization, L.P.; supervision, L.P.; project administration, L.P.; funding acquisition, L.P. All authors have read and agreed to the published version of the manuscript.

## Data availability

The unprocessed genome sequencing data have been deposited in the NCBI database under the BioSample accession SAMN18815149, within the BioProject PRJNA723377. The corresponding genome assembly is accessible through the NCBI Whole Genome Shotgun (WGS) database under the accession number JAGTPU000000000.

## Acknowledgements

Leopoldo Palma gratefully acknowledges the Spanish Ministry of Science, Innovation, and Universities, the Spanish State Research Agency, and the European Union for funding his Ramón y Cajal contract (grant ref. RYC2023-043507-I).

## Appendix A. Supplementary data

Supplementary data associated with this article, including the TYGS taxonomic report, RAST annotation tables and Swiss-Prot BLAST results are available at Zenodo (DOI: 10.5281/zenodo.19456917).

